# The interaction between genotype and maternal nutritional environments affects tomato seed and seedling quality

**DOI:** 10.1101/458836

**Authors:** Nafiseh Geshnizjani, Saadat Sarikhani Khorami, Leo A.J. Willems, Basten L. Snoek, Henk W.M. Hilhorst, Wilco Ligterink

## Abstract

Seed and seedling traits are affected by the conditions of the maternal environment, such as light, temperature and nutrient availability. In this study, we have investigated whether different maternally applied nitrate and phosphate concentrations affect the seed and seedling performance of two tomato genotypes: *Solanum lycopersicum* cv. Money maker and *Solanum pimpinellifolium* accession CGN14498. We observed large differences for seed and seedling traits between the two genotypes. Additionally, we have shown that for nitrate most of the seed and seedling traits were significantly affected by genotype by environment interactions (G×E). The effect of the maternal environment was clearly visible in the primary metabolites of the dry seeds. For example, we could show that the amount of γ-aminobutyric acid (GABA) in Money maker seeds was affected by the differences in the maternal environments and was positively correlated with seed germination under high temperature. Overall, compared to phosphate, nitrate had a larger effect on seed and seedling performance in tomato. In general, the different responses to the maternal environments of the two tomato genotypes show a major role of genotype by environment interactions in shaping seed and seedling traits.

**Highlight:** The presented data specifically provides knowledge towards understanding a multi-level effect of the maternal nutritional environment on seed and seedling characteristics in tomato. We show a clear genotype by environment interactions (G×E) especially for maternal growth on different nitrate concentrations. Additionally we identified metabolites with either positive or negative correlations with maternal environment affected phenotypical traits.

## Introduction

Seeds, as the start point of the life cycle of plants, can be considered as the key life stage in many crops like tomato. High quality and well developed seeds are crucial for a successful life cycle of crops, from seedling establishment through to fruit and seed production, especially under stressful environmental conditions. Seed quality is a complex trait and is composed of different quality characteristics including physical, physiological, genetic and seed health quality (Sperling *et al.*, 2004). In addition, seed quality is influenced by many environmental cues such as drought, light and temperature (Rowse and Finch-Savage, 2003). Establishment of seed quality starts at the position where the plants grow, produce and mature their seeds (Delouche and Baskin, 1971). The maternal environment under which seeds develop and mature, including the climate and growth conditions, has a profound influence on seed quality (Delouche, 1980). Maternal environmental effects are defined as a specific phenomenon in which the phenotype of offspring is influenced by the environment that the maternal plant is exposed to (Donohue, 2009). It has been reported that different temperatures (Demir *et al.*, 2004; He *et al.*, 2014; Schmuths *et al.*, 2006), photoperiod (Munir *et al.*, 2001; Pourrat and Jacques, 1975) and nutrient conditions (Alboresi *et al.*, 2005; He *et al.*, 2014) during seed development and maturation may result in differences in seed performance in plants such as tomato and Arabidopsis.

Seed performance traits, such as seed dormancy and germinability, can be influenced by different environmental conditions. The germinability of a seed batch is defined as the percentage of seed germination during a specific time interval (Fenner, 1991). There are many reports on the influence of environmental conditions under which seeds develop and mature on seed dormancy and germinability. For instance, for *Solanum lycopersicum* (Varis and George, 1985), *Nicotiana tabacum* (Thomas and Raper, 1979), *Sisymbrium officinale* (Hilhorst and Karssen, 1988), *Arabidopsis thaliana* (Alboresi *et al.*, 2005; He *et al.*, 2014) and *Rumex crispus* (Hejcman *et al.*, 2012) it has been shown that low nitrate levels in the soil of the mother plant results in a decrease in germinability of their seeds. Alboresi et al. (2005) reported that nitrate can reduce dormancy in Arabidopsis seeds by either direct effects or through hormonal and metabolic changes in the seed. These changes probably include interactions with ABA and/or GA synthesis and degradation pathways (Alboresi *et al.*, 2005). The effects of maternal environmental conditions on seed quality are not restricted to germination characteristics of the seeds, but may also include other seed quality traits such as seed size and seed weight as well as seedling quality characteristics such as root and shoot weight, hypocotyl length and root architecture. In many species a higher level of nutrient supply to the mother plant led to the production of bigger and heavier seeds (Fenner, 1992). Moreover, in some species a higher nitrate regime applied to the mother plant resulted in an enhanced seedling establishment and higher shoot and root weight of the seedlings (Farhadi *et al.*, 2014; Song *et al.*, 2014). In addition, there are many examples of changed metabolism in seeds in response to the environmental condition of the mother plant (Joosen *et al.*, 2013; Mounet *et al.*, 2007). A better understanding of the influence of the maternal environment on seed and seedling quality can be obtained by performing omics analysis of seeds such as transcriptomics, proteomics and metabolomics.

Fait *et al.* (2006) revealed that seed germination and seedling establishment characteristics are associated with degradation and mobilization of reserves which are accumulated during seed maturation like sugars, organic acids and amino acids. Therefore, profiling the metabolites and finding the ones associated with phenotypes can be regarded as a powerful tool for monitoring seed performance. In general, metabolite contents alter in response to abiotic stress, which is most obvious for primary metabolites such as sugars, amino acids and tricarboxylic acid (TCA) cycle intermediates (Arbona *et al.*, 2013).

In this study, we investigated if different maternal nutritional environments can affect the quality of the progeny of different genotypes. For this purpose we investigated two different tomato genotypes (*Solanum lycopersicum* cv. Money maker (MM) and *Solanum pimpinellifolium* (PI)) under different nutrient conditions.

PI, the most closely related wild tomato species to the advanced tomato breeding line (MM), has been used in breeding programs for its tolerance to some sub-optimal environments as well as the ability of being naturally crossed with this species. We grew these genotypes in different concentrations of nitrate and phosphate. Phosphate is an important nutrient for plants, making up 0.2% of the dry weight and being an essential part of some vital molecules like nucleic acids, phospholipids and ATP. Nitrate plays a key role in plants as a major source of nitrogen and some signal metabolites (Schachtman *et al.*, 1998; Urbanczyk-Wochniak and Fernie, 2005). Under both optimal and stressful conditions extensive phenotyping by germination tests and metabolite profiling was done after harvesting the seeds. Based on these results we show that different levels of phosphate and nitrate available to the mother plant can influence seed and seedling traits especially under stressful germination conditions. In addition, in order to investigate if physiological changes in seed and seedling performance are influenced by metabolic changes in the dry seed, correlation analysis was performed between physiological traits like seed germination and seedling growth and metabolic changes caused by the different maternal environments in tomato. We showed that several phenotypic traits are either positively or negatively correlated with metabolites.

## Materials and Methods

### Plant material, growth condition and seed extraction

MM and PI plants were grown under standard nutrient conditions (table 1, Supplementary Table S1) with a 16-h light and 8-h dark photoperiod. The temperature was controlled during the day and night at 25°C and 15°C, respectively. From first open flower onwards the plants were transferred to the different nutrient conditions (table 1, Supplementary Table S1). For each environment four biological replicates were used. All plants were grown in the greenhouse at Wageningen University, the Netherlands. After harvesting, the seeds were collected from healthy and ripe fruits. In order to remove the pulp attached to the seeds, they were treated with 1% hydrochloric acid (HCl) and subsequently passed through a mesh sieve and washed with water to remove the remaining HCl and pulp. In the following step, seeds were treated with trisodium phosphate (Na3PO4.12H2O) for disinfection. Finally, seeds were dried at 20°C for 3 days on a clean filter paper in ambient conditions and stored in the paper bags at the room temperature (Kazmi *et al.*, 2012).

**Table 1.**
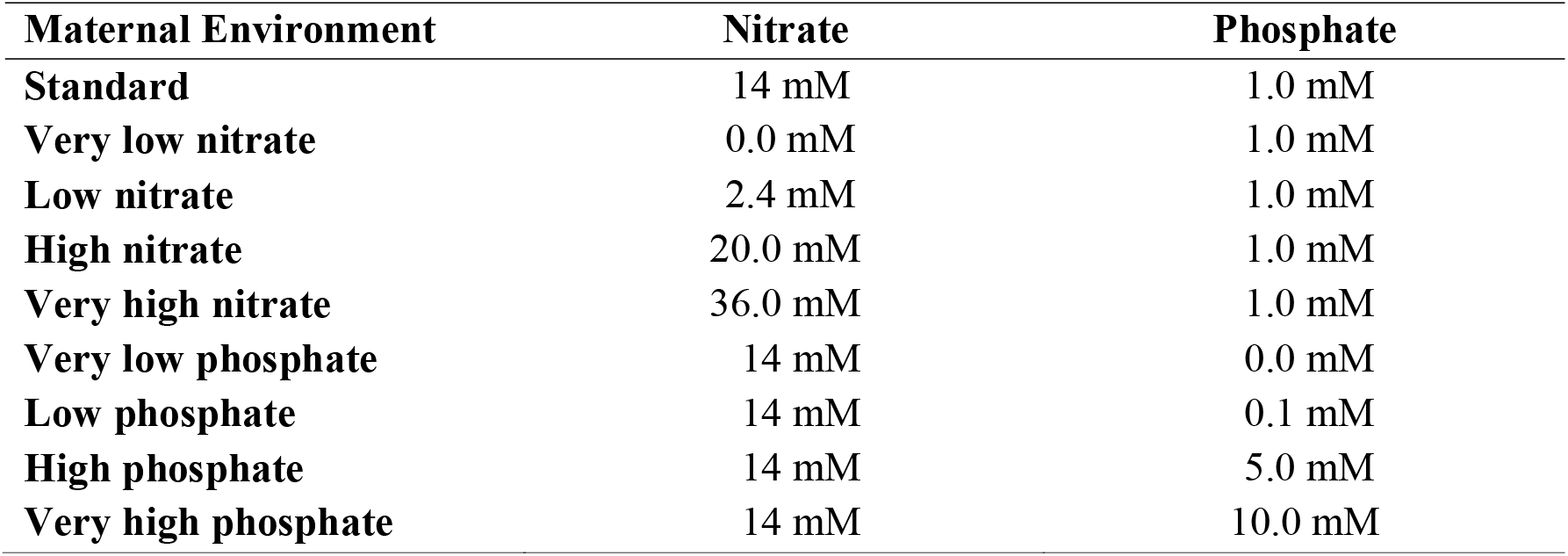
Nutrient conditions of mother plants after flowering.

| **Maternal Environment** | **Nitrate** | **Phosphate** |
| --- | --- | --- |
| **Standard** | 14 mM | 1.0 mM |
| **Very low nitrate** | 0.0 mM | 1.0 mM |
| **Low nitrate** | 2.4 mM | 1.0 mM |
| **High nitrate** | 20.0 mM | 1.0 mM |
| **Very high nitrate** | 36.0 mM | 1.0 mM |
| **Very low phosphate** | 14 mM | 0.0 mM |
| **Low phosphate** | 14 mM | 0.1 mM |
| **High phosphate** | 14 mM | 5.0 mM |
| **Very high phosphate** | 14 mM | 10.0 mM |

### Seed phenotyping

#### Seed size and weight

Seed size was determined by taking photographs of 12-h imbibed seeds on white filter paper (20.2 x 14.3 cm) using a Nikon D80 camera fixed to a repro stand with 60 mm objective and connected to a computer with Nikon camera control pro software version 2.0 (Joosen *et al.*, 2010). The pictures were analysed by ImageJ (http://rsbweb.nih.gov/ij/) combining colour threshold with particle analysis. Seed weight was measured by weighing approximately 100 dry seeds and divided by the number of seeds.

#### Germination assay

Germination assays were performed with four replications of around 50 seeds per sample of both genotypes in a completely randomized design. The seeds were sown in germination trays (21×15 cm DBP Plastics, http://www.dbp.be) on two blue germination papers (5.6′ x 8′ Blue Blotter Paper; Anchor Paper Company, http://www.seedpaper.com) and 50 ml demineralized water in the case of optimal and high temperature germination environments or Sodium Chloride (−0.5 MPa NaCl; Sigma-Aldrich) and mannitol (−0.5 MPa; Sigma-Aldrich) in the salt and osmotic stress conditions, respectively. Each germination tray contained three samples, using a special mask to ensure correct placement. The trays were piled up in different piles with two empty trays on the top and bottom, containing two white filter papers and 15ml of water and covered by white plastic lids to prevent unequal evaporation and wrapped in a transparent plastic bag and stored at 4°C for 3 days. Subsequently, the bags were transferred to an incubator (type 5042; seed processing Holland, http://www.seedprocessing.nl) in the dark at 25°C except for high temperature which was at 35°C. Germination was scored at 24-h intervals during 14 consecutive days in the case of salt and osmotic stress conditions and at 8- h intervals for one week for the optimal and high temperature conditions.

### Seedling phenotyping

Seedling characteristics were measured in two separate experiments. In the first 12 x 12 cm petri dishes, filled with half MS medium with agar (1%) were used. The top 4 cm of the medium was removed and the seeds, which were sterilized for 16 h in a desiccator above 100 ml sodium hypochlorite (4%) with 3 ml concentrated HCl, were sown on top of the remaining 8 cm. After sowing the seeds, the plates were stored in the cold room (4°C) for 3 days and subsequently transferred to a climate chamber and held in a vertical position (70° angle) under 25°C with 16h light and 8h dark. For each plate 14 seeds were used and the first 7 germinated seeds were kept. Germination was scored during the day at 8-h intervals as visible radical protrusion. After the start of germination pictures were taken at 24-h intervals for root architecture analysis. Five days after germination, seedlings were harvested and hypocotyl length (HypL) was measured. EZ-Rhizo was used to analyse root architecture (Armengaud *et al.*, 2009) and main root length (MRL) and number of lateral roots (NLR) were determined. In the second experiment, 20 seeds of each seed batch were sown in germination trays and stored for 3 days at 4°C. Afterwards they were transferred to an incubator at 25°C. The first 10 germinated seeds were placed on round blue filter papers (9 cm Blue Blotter Paper; Anchor Paper Company, http://www.seedpaper.com) on a Copenhagen table at 25°C in a randomized complete block design (with 4 biological replicates) for 10 days. To prevent evaporation, conical plastic covers with a small hole on top were placed on top of the filter papers. After 10 days, fresh and dry weight of root and shoot of the seedlings was measured (FWR, DWR, FWSH and DWSH respectively).

### Nitrate, phosphate and phytate measurement

To determine the nitrate, phosphate and phytate content of the seed samples, 15-20 mg of dry seeds were frozen in liquid nitrogen and homogenized in a dismembrator (Mikro-dismembrator U; B. Braun Biotech International, Melsungen, Germany), by using 0.6 cm glass beads, at 2500rpm for 1 minute. Fifteen mg of dry homogenized seeds with 1 ml 0.5 N HCl and 50 mg l^-1^ *trans-aconitate* (internal standard) was incubated at 100°C for 15 minutes. After centrifugation for 3 minutes at 14000 rpm, the supernatant was filtered using Minisart SRP4 filters (Sartorius Stedim Biotech, http://www.sartorius.com) and transferred to an HPLC-vial.

A Dionex ICS2500 system was used for HPLC-analysis with an AS11-HC column and an AG11-HC guard column. The elution was performed by 0–15 min linear gradient of 25–100 mM NaOH followed by 15–20 min 500 mM NaOH and 20–35 min 5 mM NaOH with a flow rate of 1 ml min^-1^ throughout the run. Contaminating anions in the samples were removed by an ion trap column (ATC) which was installed between the pump and the sample injection valve. Conductivity detection chromatography was performed for anion detection, an ASRS suppressor was used to reduce background conductivity and water was used as counter flow. Identification and quantification of peaks was done by using authenticated external standards of nitrate (NaNO3, Merck), phosphate (Na2HPO4.2H2O, Merck) and phytate (Na(12)-IP6 IP6, Sigma-Aldrich).

### ABA determination

For ABA determination, approximately 15 mg of dry weight seed samples were homogenized as described above for nitrate, phosphate and phytate extraction and extracted in 1 ml of 10% methanol/water (v/v) according to Floková et al. (Floková *et al.*, 2014) with modifications. Stable isotope-labelled internal standard of [2H6]-ABA was added to each sample in order to validate ABA quantification. Sample extracts were centrifuged (13000 rpm/10 min/4°C) and further purified by solid-phase extraction using Strata X (30 mg/3 cc, Phenomenex) columns, activated with 1 ml of methanol, water and 1 ml of the extraction solvent. The loaded samples were washed with 3 ml of water and analyte elution was performed with 3 ml of 80% methanol/water (v/v). Samples were evaporated to dryness in a Speed-Vac concentrator and reconstituted in 60 μl of mobile phase prior to the UPLC-MS/MS analysis. The Acquity UPLC^®^ System (Waters, Milford, MA, USA) coupled to a triple quadrupole mass spectrometer Xevo™ TQ S (Waters MS Technologies, Manchester, UK) was employed to measure ABA levels. Samples were injected on a reverse phase based column Acquity UPLC^®^ CSH^™^ C18; 2.1 x 100 mm; 1.7 μm (Waters, Ireland) at flow rate 0.4 ml min^-1^. Separation was achieved at 40°C by 9 min of gradient elution using A) 15 mM formic acid/water and B) acetonitrile: 0-1 min isocratic elution at 15% B (v/v), a 1-7 min linear gradient to 60% B, 7-9 min linear gradient to 80% B and a 9-10 min logarithmic gradient to 100% B. Finally, the column was washed with 100% acetonitrile and equilibrated to initial conditions for 2 min. the eluate was introduced to the electrospray ion source of tandem mass spectrometer operating at the following settings: source/desolvation temperature (120/550°C), cone/desolvation gas flow (147/650 l h^-1^), capillary voltage (3 kV), cone voltage (30 V), collision energy (20 eV) and collision gas flow 0.25 ml min^-1^. ABA was quantified in multiple reaction monitoring mode (MRM) using standard isotope dilution method. The MassLynx™ software (version 4.1, Waters, Milford, MA, USA) was used to control the instrument, MS data acquisition and processing.

### Analysis of seed metabolites by GC-TOF-MS

For metabolite extraction we used the method as described by Roessner *et al.* (2000)with small modifications. Approximately 30 tomato seeds were homogenized with a micro dismembrator (Sartorius) in 2 ml Eppendorf tubes with 2 iron beads (2.5 mm) precooled with liquid nitrogen and then 10 mg of that material has been used for metabolite extraction. Metabolite extraction was done by adding 700 μl methanol/chloroform (4:3) together with a standard (0.2 mg/ml ribitol) to each sample and mixed thoroughly. Samples were sonicated for 10 minutes and 200 μl Mili-Q water was added, followed by vortexing and centrifugation (5 min, 13500 rpm). The methanol phase was collected and transferred to a new 2 ml Eppendorf tube. Five hundred μl methanol/chloroform was added to the remaining organic phase, kept on ice for 10 min followed by adding 200 μl Mili-Q water. After vortexing and centrifugation (5 min, 13500 rpm), the methanol phase was collected and added to the previous collected phase. Finally, 100 μl of total extract was transferred to a glass vial and dried overnight in a speedvac centrifuge at 35°C (Savant SPD1211).

For each maternal environment four biological replicates were used and the gas chromatography-time of flight-mass spectrometry (GC-TOF-MS) method was used for metabolite analysis which was previously described by Carreno-Quintero *et al.* (2012). Detector voltage was set at 1600 V. The chromaTOF software 2.0 (Leco instruments) was used for analysing the raw data and further processing for extracting and aligning the mass signals was performed using the Metalign software (Lommen, 2009). A signal to noise ratio of 2 was used. Afterwards, the output was further analysed using the Metalign output Transformer (METOT; Plant Research International, Wageningen) and Centrotypes were constructed using MSClust (Tikunov *et al.*, 2012). The identification of Centrotypes was performed by matching the mass spectra to an in-house-constructed library, to the GOLM metabolome database (http://gmd.mpimp-golm.mpg.de/) and to the NIST05 library (National Institute of Standards and Technology, Gaithersburg, MD, USA; http://www.nist.gov/srd/mslist.htm). The identification was based on similarity of spectra and comparison of retention indices calculated using a 3^th^ order polynomial function (Strehmel *et al.*, 2008).

### Statistical analysis

#### Calculation of G_max_, t_50_^-1^, AUC and U_8416_

Seed performance was determined by calculating maximum germination (G_max_, %), rate of germination or the reciprocal of time to reach 50% of germination (t_50_^-1^), uniformity of germination or time from 16% till 84% germination (U_8416_, h) and area under the germination curve (AUC, during the first 100 and 200h for optimal and high temperature respectively and 300h in the case of salt and mannitol stress conditions) using the curve-fitting module of the Germinator package (Joosen *et al.*, 2010, Ligterink and Hilhorst, 2017).

#### Analysis of all factors affecting seed and seedling traits: Genotype, Environments and Genotype by environment interactions (G×E)

To identify the factors correlating with seed and seedling traits we used an ANOVA (with linear model trait ~ genotype * treatment). The different treatment regimens (N and P) where studied separately. A significance threshold of 0.05 was used.

#### Cluster, principle component analysis and correlation analysis

Cluster and principle component analysis (PCA) were performed using the online web tool MetaboAnalyst 3.0; www.metaboanalyst.ca (Xia *et al.*, 2015).

R-packages “MASS”, “Hmisc”, “VGAM”, “gplots” and “graphics” (https://www.r-project.org/) were used for analysis and construction of the correlation between measured traits.

## Results

Several studies have been reported recently about the effect of the maternal environment such as temperature, light and nutrition on seed and seedling quality in plants (Alboresi *et al.*, 2005; He *et al.*, 2014; Hejcman *et al.*, 2012). However there is still a lack of knowledge on the influence of nutritional condition of the mother plant on seed and seedling performance. In order to investigate the effect of maternal nutrient environment on the seed and seedling quality in tomato, two tomato genotypes, MM and PI were grown on different nutrient solutions from flowering onwards (Table 1). Their seeds were harvested and phenotyped for various seed performance traits, including percentage of germination, germination rate and uniformity, under optimal and several stress germination conditions (i.e. high temperature, salt and osmotic stress). Furthermore, seed size and weight were determined. Since final successful and sustainable crop production results from healthy seedlings and good seedling establishment, we also measured some seedling quality traits such as hypocotyl length, root architecture and fresh and dry root and shoot weight.

### Factors affecting seed and seedling traits

A linear model/ANOVA was used to investigate the effects caused by the different factors like genotype, environment and their interaction (G×E). The results showed that genotype was an important factor, since it had a very pronounced influence on almost all traits in different nitrate and phosphate concentrations (Table 2). Also the environment under which seeds developed had a significant effect which was most prominent for different nitrate conditions (Table 2). Although G×E interactions had a significant effect on some traits for the phosphate environment, most of the seed and seedling traits in the case of different nitrate concentrations were significantly influenced by G×E interactions (Table 2).

**Table 2.** ANOVA analysis of the effect of genotype, maternal environment and genotype-by-environment interactions on seed and seedling quality. Values show the -10 log(*P*) ^*^.

| Traits | Nitrate |  |  | Phosphate |  |  |
| --- | --- | --- | --- | --- | --- | --- |
|  | Genotype | Environment | G×E | Genotype | Environment | G×E |
| Seed Size | 25.24 | 7.81 | 4.12 | 23.37 | 2.41 | 0.47 |
| Seed Weight | 27.38 | 8.88 | 6.06 | 24.17 | 5.71 | 3.91 |
| FWR | 22.90 | 3.29 | 1.88 | 18.22 | 3.64 | 1.29 |
| DWR | 22.42 | 4.09 | 3.85 | 16.60 | 3.16 | 0.75 |
| FWSH | 25.07 | 5.89 | 3.19 | 18.45 | 3.64 | 2.32 |
| DWSH | 22.91 | 4.67 | 2.95 | 20.05 | 5.07 | 2.54 |
| MRL | 16.23 | 4.29 | 1.31 | 14.61 | 0.59 | 0.83 |
| NLR | 16.08 | 0.04 | 0.43 | 17.05 | 0.48 | 1.49 |
| G<sub>max</sub> Water | 5.74 | 3.18 | 3.05 | 5.59 | 0.32 | 0.14 |
| t<sub>50</sub><sup>-1</sup> Water | 16.72 | 0.26 | 0.20 | 23.78 | 0.06 | 0.93 |
| G<sub>max</sub> NaCl | 13.57 | 2.25 | 2.31 | 10.16 | 1.55 | 1.04 |
| t<sub>50</sub><sup>-1</sup> NaCl | 16.85 | 0.11 | 1.02 | 27.48 | 1.89 | 0.92 |
| G<sub>max</sub> HT | 27.43 | 3.45 | 3.63 | 24.73 | 1.88 | 2.27 |
| t<sub>50</sub><sup>-1</sup> HT | 29.69 | 11.88 | 4.08 | 23.38 | 0.89 | 1.58 |
| G<sub>max</sub> Mannitol | 16.27 | 7.77 | 8.79 | 9.98 | 0.02 | 0.01 |
| t<sub>50</sub><sup>-1</sup> Mannitol | 23.71 | 7.55 | 10.62 | 16.50 | 0.71 | 2.14 |
| Nitrate Content | 1.66 | 0.06 | 0.77 | 2.15 | 3.03 | 3.39 |
| Phytate Content | 12.06 | 2.74 | 1.90 | 16.85 | 21.16 | 4.22 |
| ABA Content | 6.84 | 1.04 | 1.04 | 4.95 | 3.21 | 1.12 |

\* Coloured cells demonstrate significant levels (Dark green: $P < 0.001$ ; Light green: $P < 0.01$ ; Very light green: $P < 0.05$ ) and non-coloured spots represent non-significant values.

### The effect of different nutrient regimes of the mother plant on seed quality traits

#### Seed germination under optimal conditions (water)

Under normal germination conditions only very low nitrate (0 mM) decreased the germination percentage in MM (Fig. 1A). Although the rate of the germination (t_50_ ^-1^) was not affected significantly by different amounts of nitrate, it was decreased by higher amounts of phosphate (Supplementary Fig. S1A).

**Fig. 1.**
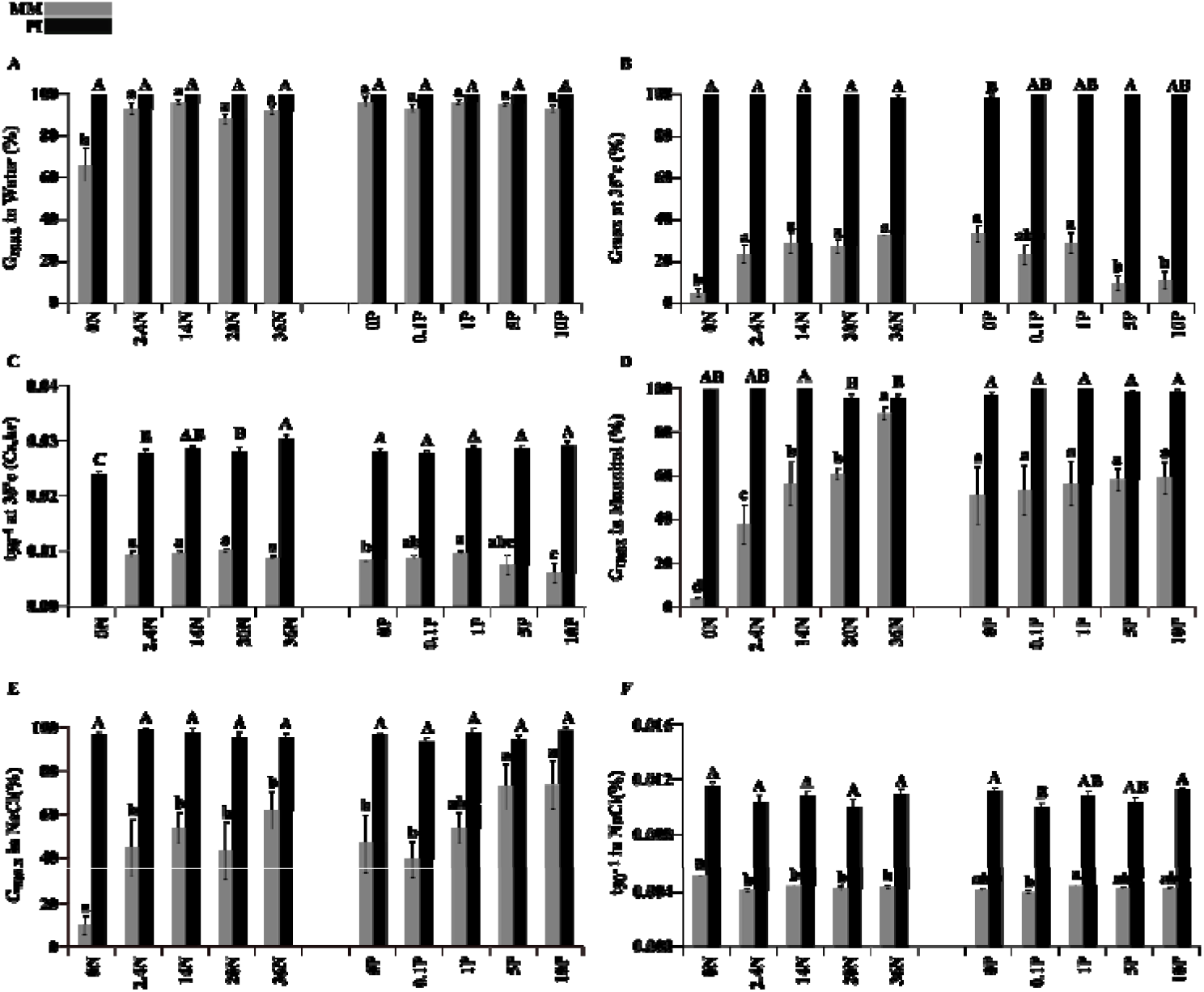
Effects of maternal nutritional environment on seed germination traits of **MM** and **PI**. **A**, Germination in water; **B**, Germination at high temperature (35°C); **C**, t_50_^-1^ at high temperature (35°C); **D**, Germination in mannitol (−0.5 MPa); **E**, Germination in salt (−0.5 MPa); **F**, t_50_^-1^ in salt (−0.5 MPa) in different concentrations of nitrate (0N, 2.4N, 14N, 20N and 36N) and phosphate (0P, 0.1P, 1P, 5P and 10P). Letters above the bars represent significant differences between different concentrations of nitrate or phosphate within each genotype (p<0.05).

#### Seed germination in stress conditions (high temperature, salt and mannitol)

Our results showed that at high temperature MM seeds from plants grown in 0 mM nitrate, germinated very poorly (4%) while higher concentrations of nitrate resulted in significantly higher germination percentages (40-60%; Fig. 1B). These seeds also had a higher t_50_^-1^ (Fig. 1C). In contrast with nitrate, G_max_ was decreased at higher levels of phosphate (Fig. 1B). While seed germination of MM was positively correlated with nitrate concentration in mannitol (Fig. 1D), germination rate was contrarily decreased at higher levels of both nutrients (Supplementary Fig. S1B). Under salt stress, both phosphate and nitrate had a positive effect on germination percentage of MM seeds and a negative effect on their germination rate (lower t50^-1^ values, Fig. 1E, F).

Although different nutritional environments resulted in clear changes in seed quality traits in MM, hardly any effect was seen for PI seeds, indicating that PI was tolerant to the different environments that were tested.

#### Seed size and weight

By increasing the nitrate level, seed size and weight of MM plants increased. However, both seed size and weight decreased slightly again at concentrations of 20 mM nitrate or higher. For PI, higher amounts of nitrate and phosphate led to the production of larger and heavier seeds (Fig. 2A,B).

**Fig. 2.**
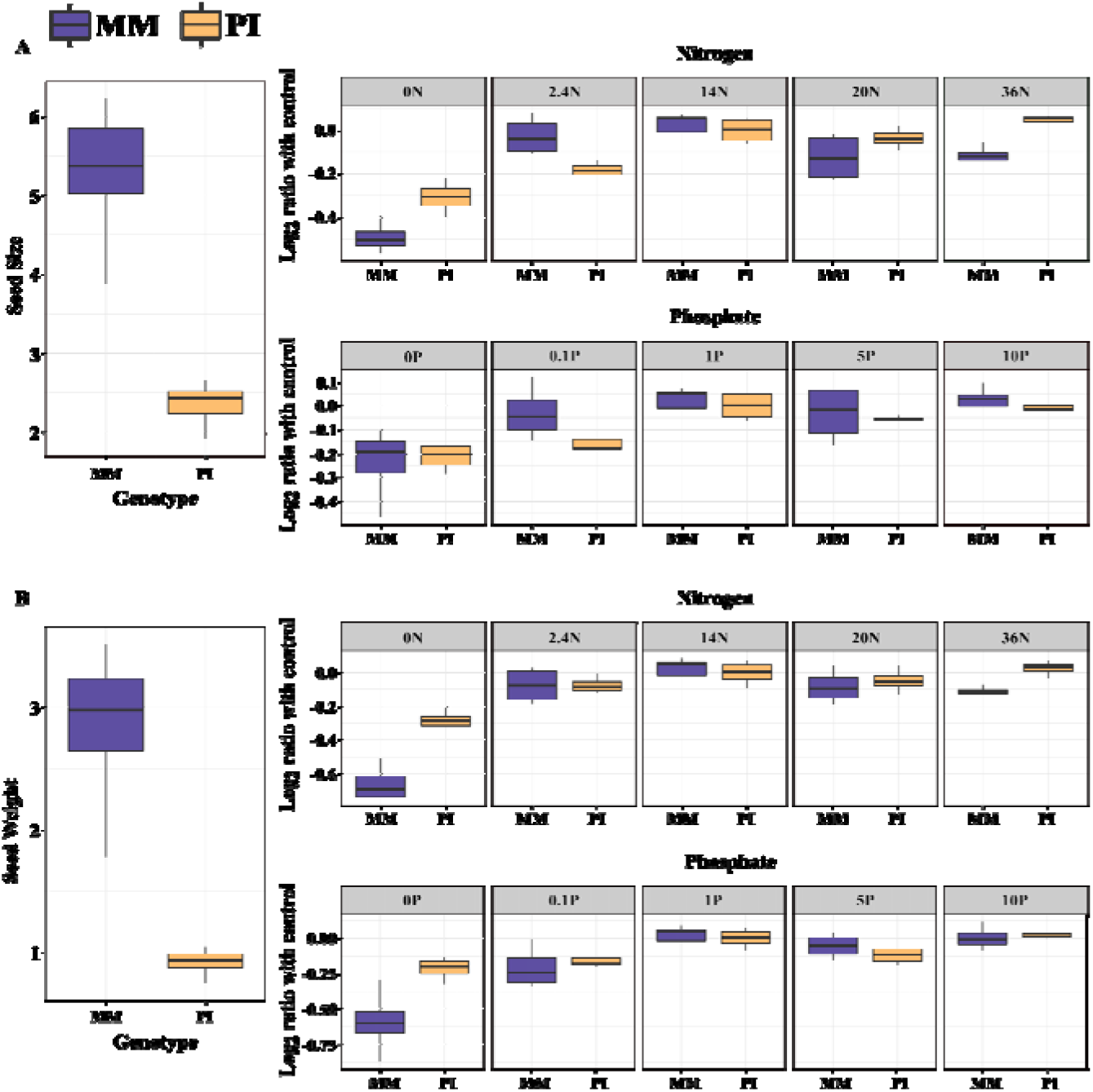
Effects of maternal nutritional environments on seed quality of **MM** and **PI**. **A**, Seed size; **B**, Seed weight of the plants grown in different concentrations of nitrate (0N, 2.4N, 14N, 20N and 36N) and phosphate (0P, 0.1P, 1P, 5P and 10P). On left, the average of seed size and seed weight (regardless of maternal environments) in each genotype are presented.

### ABA, nitrate and phytate

ABA content of dry seeds was not significantly influenced by the maternal nitrate concentration, but was increased by application of 1 mM of phosphate. Although ABA showed a relatively consistent increase in PI, concentrations above 1 mM of phosphate resulted in decreased ABA levels in MM seeds (Supplementary Fig. S2A). The phytate content of the seeds significantly increased with higher phosphate levels in both genotypes (Supplementary Fig. S2B). Application of nitrate up to 14 mM increased phytate levels of MM seeds. However, concentrations above 14 mM led to decreased phytate levels in both genotypes (Supplementary Fig. S2B). In PI seeds nitrate content was not affected by the nutrient nitrate level, while in MM higher levels of nitrate surprisingly led to lower seed nitrate levels.

### The effects of different nutrient regimes of the mother plant on seedling quality traits

#### Fresh and dry shoot and root weight

Both FWR and FWSH of seedlings were influenced by different concentrations of nitrate and phosphate for the mother plant. Evidently, raising the dosage of nitrate and phosphate in both MM and PI resulted in heavier seedlings (shoot and root) (Fig. 3A, B). Shoot and root dry weight followed the same pattern as that of the fresh weight in different environments in both lines (Supplementary Fig. S3A, B).

**Fig. 3.**
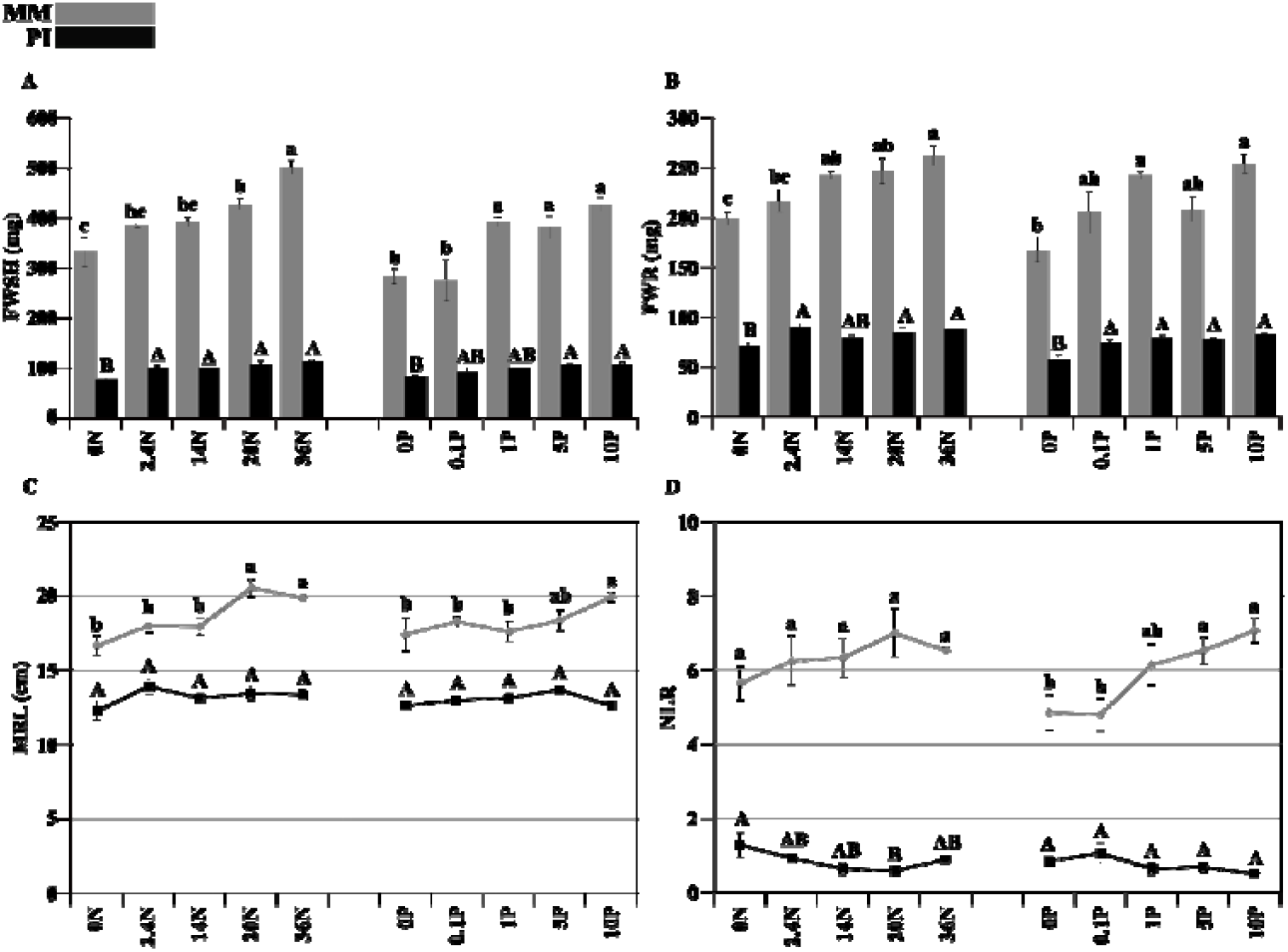
Effects of maternal nutritional environments on seedling quality traits of **MM** and **PI**. **A**, Shoot fresh weight; **B**, Root fresh weight; **C**, Main root length; **D**, Number of lateral roots in different concentrations of nitrate (0N, 2.4N, 14N, 20N and 36N) and phosphate (0P, 0.1P, 1P, 5P and 10P). Letters above the bars (A, B) and lines (C, D) represent significant differences between different concentrations of nitrate or phosphate within each genotype (p<0.05).

#### Root architecture

Although higher amounts of nitrate and phosphate produced a lower NLR in PI, MRL of these plants were not remarkably influenced by different nutritional environments (Fig. 3C, D). In contrast, MM plants grown in higher regimes of nitrate and phosphate produced seedlings with longer main roots and a higher NLR (Fig. 3C, D). Hypocotyl length of the seedlings was not influenced significantly by the maternal environment (Supplementary Fig. S3C).

### Trait by trait correlation

In order to investigate how different maternal nutrient environments affected different seed and seedling characteristics in a similar way, a correlation analysis was performed for all pairs of measured traits for either different concentrations of nitrate or phosphate, separately (Fig. 4, Supplementary Table S1). For the nitrate environment, nitrate levels were positively correlated with seed and seedling performance traits such as seed size, seed weight and FWSH and FWR, however nitrate content of the seeds was negatively correlated with nutrient nitrate levels (Fig. 4). ABA levels had a negative correlation with almost all the measured phenotypes as also has been observed for *A. thaliana* (He *et al.*, 2016). For the phosphate environment, seed size, seed weight, germination in mannitol and salt, FWR, FWSH and phytate content were strongly correlated with phosphate levels. Moreover seed size and seed weight also showed a strong positive correlation with FWR and FWSH of seedlings for the different phosphate environments (Fig. 4).

**Fig. 4.**
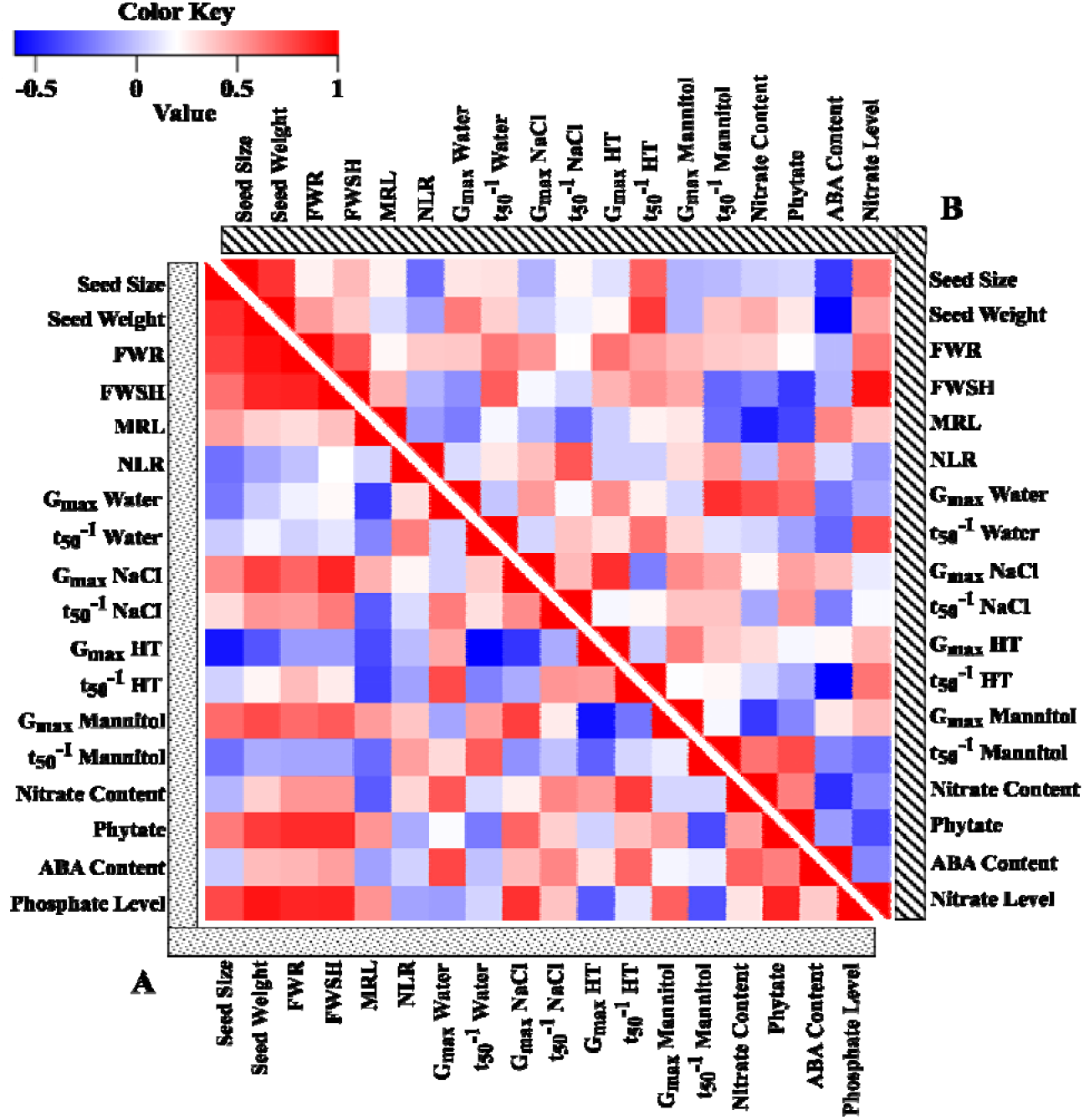
Heatmap of trait by trait correlations of seed and seedling traits in **MM** and **PI**: in response to different concentration of **(A)** phosphate and **(B)** nitrate.

### Metabolite analysis

Nutritional environments of the mother plant affected seed and seedling performance traits. Since the metabolites in the dry seeds have been built up during seed maturation and drying, the underlying metabolic pathways have been analysed using a metabolomics approach to see if the observed differences in phenotype can be explained by different metabolic content of the mature dry seeds. Dry mature seeds from plants grown in the different nutritional environments have been used for metabolic analysis as it gives a broad overview of the biochemical status of the seeds and helps to better understand the responses to the different environments. This resulted in the detection of 89 primary metabolites from which 50 could be identified. These could be classified as amino acids, organic acids, sugars and some other metabolites which are intermediates of key metabolic compounds (Supplementary Excel File S1). MM plants grown with 0 mM nitrate produced less seeds which have been used for the germination assays and therefore, metabolites of these seeds could not be measured in this study.

#### Genetic effects on metabolite profiles

Both PCA and cluster analysis of the metabolites showed that metabolite content was mainly affected by the genetic background of the seeds. A clear separation between samples of the two genotypes in terms of known metabolites was observed in a PCA plot which indicated that the metabolic variation caused by genetic background was larger than the variation caused by the maternal environment (Fig. 5). The dendrogram which was created by cluster analysis revealed an obvious segregation between the two genotypes which is already shown by PCA analysis. There are three main clusters for each genotype in which P-Control, P-5P, P-10P and P-2.4N; P-20N and P-36N; and P-0P, P-0.1P and P-0N were grouped together (Supplementary Fig. S4). Different environments were clustered with an almost identical pattern for MM seeds (Supplementary Fig. S4).

**Fig. 5.**
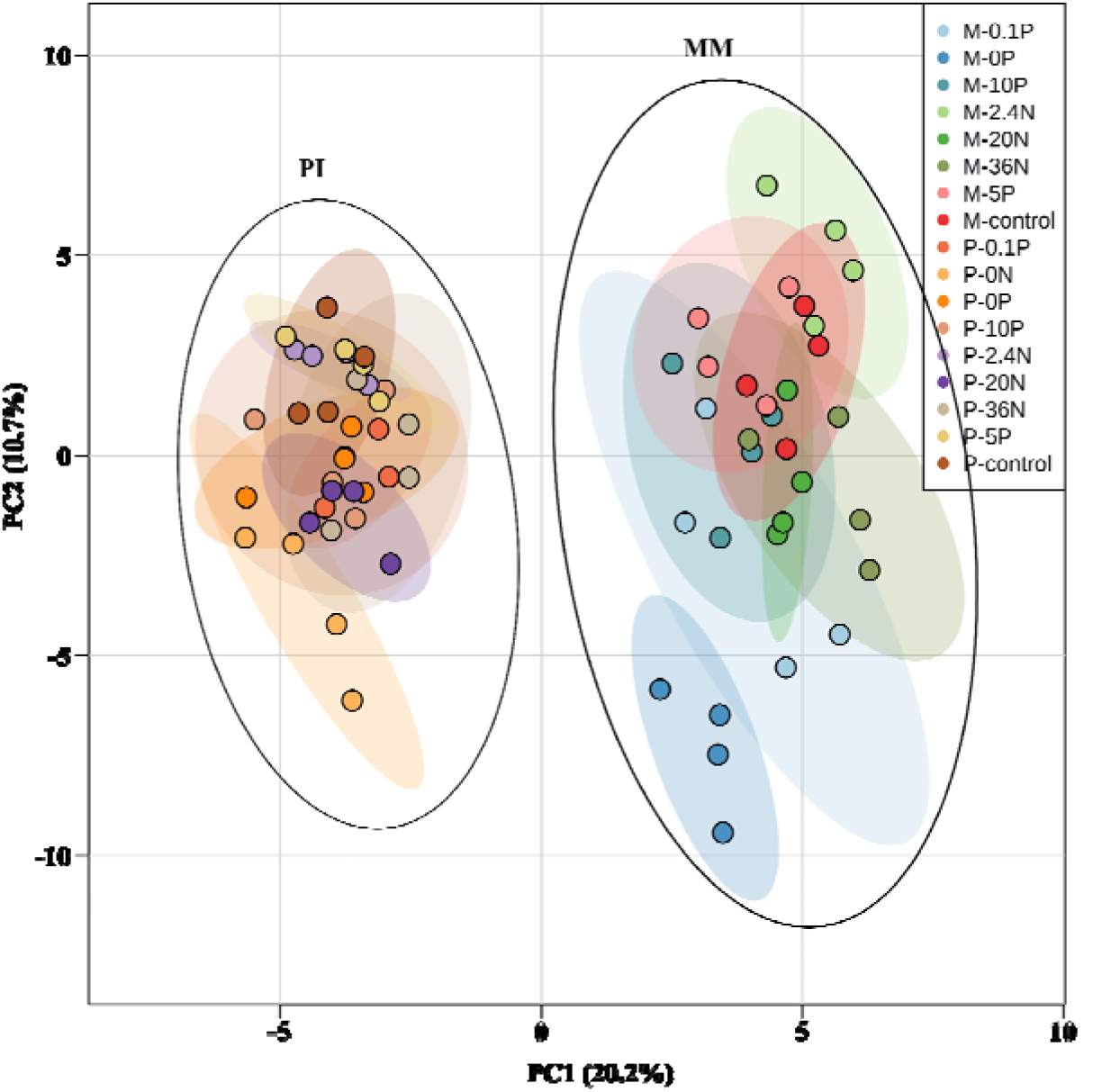
Principle component analysis of known primary metabolites in **MM, (M)** and **PI, (P)** seeds in response to different concentration of nitrate (0N, 2.4N, 14N, 20N and 36N) and phosphate (0P, 0.1P, 1P, 5P and 10P) during maternal growth.

#### Metabolic changes in response to the maternal nutrient levels

From the 50 identified metabolites, 46 were successfully mapped to their representative pathways with help of Mapman (http://MapMan.gabipd.org) and this was used to generate a metabolic framework (Fig. 6). Changing metabolite contents within the genotypes and different nutritional environments are displayed as heatmap plots below the metabolites which significantly changed in response to at least one environmental factor (Fig. 6). In general, contents of nitrogen-metabolism related metabolites such as amino acids (asparagine, alanine and γ-aminobutyric acid (GABA)) and urea were decreased significantly in seeds from plants grown under lower amounts of nitrate for both genotypes. The GABA content of MM seeds was decreased at higher levels of phosphate while it was increased at higher nitrate levels. Galactarate and pyroglutamate which both are precursors of glutamate were also increased by higher amounts of nitrate. Furthermore, some of the glycolysis and TCA cycle intermediates were remarkably affected by the maternal environment. Fructose-6-phosphate (F6P), which is one of the derivatives of glucose in the glycolytic pathway, was reduced by higher phosphate levels. Citrate and malate are two TCA intermediates which were negatively influenced by increasing phosphate levels (Fig. 6).

**Fig. 6.**
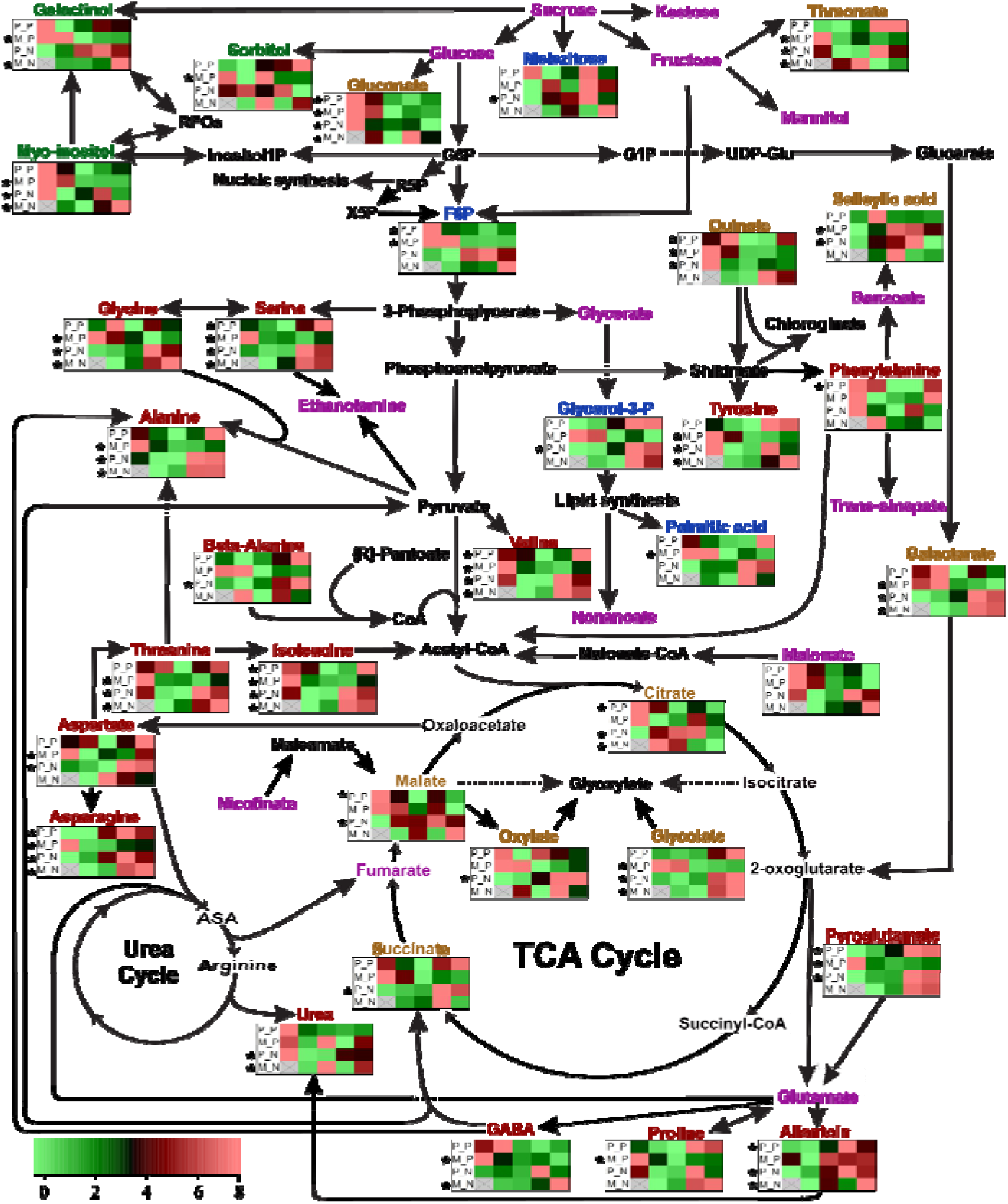
Overview of metabolic changes between the genotypes influenced by maternal nutritional environments. Metabolites are shown in three colours: **Black**, Non detected metabolites; **Purple**, Detected metabolites not significantly influenced by environment; Other colours, Detected metabolites, in different categories, significantly influenced by at least one environment; **Red**, Amino acid; **Light Brown**, Organic acid; **Green**, Sugars and sugar alcohols; **Blue**, Other categories. Heatmaps contain four rows: top two rows represent PI **(P-P)** and MM **(M-P)** in different concentrations of phosphate (0, 0.1, 1, 5 and 10 mM from left to right). The bottom two rows represent PI **(P-N)** and MM **(M-N)** in different concentrations of nitrate (0, 2.4, 14, 20 and 36 mM). Colour key represents the normalized metabolite content of seeds.

#### Correlation of seed and seedling quality traits with metabolites

A correlation analysis was performed to find correlations between metabolic changes and seed and seedling performance. In the 9 plots of Figure 7 each plot represents the correlation of metabolites with one specific trait shown in four rows (MM and PI in nitrate and MM and PI in phosphate). Correlation analysis showed that there are some traits which have similar correlation patterns for all four conditions. For example, phytate content is positively correlated with germination characteristics under saline conditions such as Gmax and t_50_ for both environmental factors in both genotypes. Furthermore there is a negative correlation of GABA and gluconate with seed size and seed weight in all four cases. There is also a positive relationship between allantoin and FWR and FWSH in all four cases (Fig. 7).

**Fig. 7.**
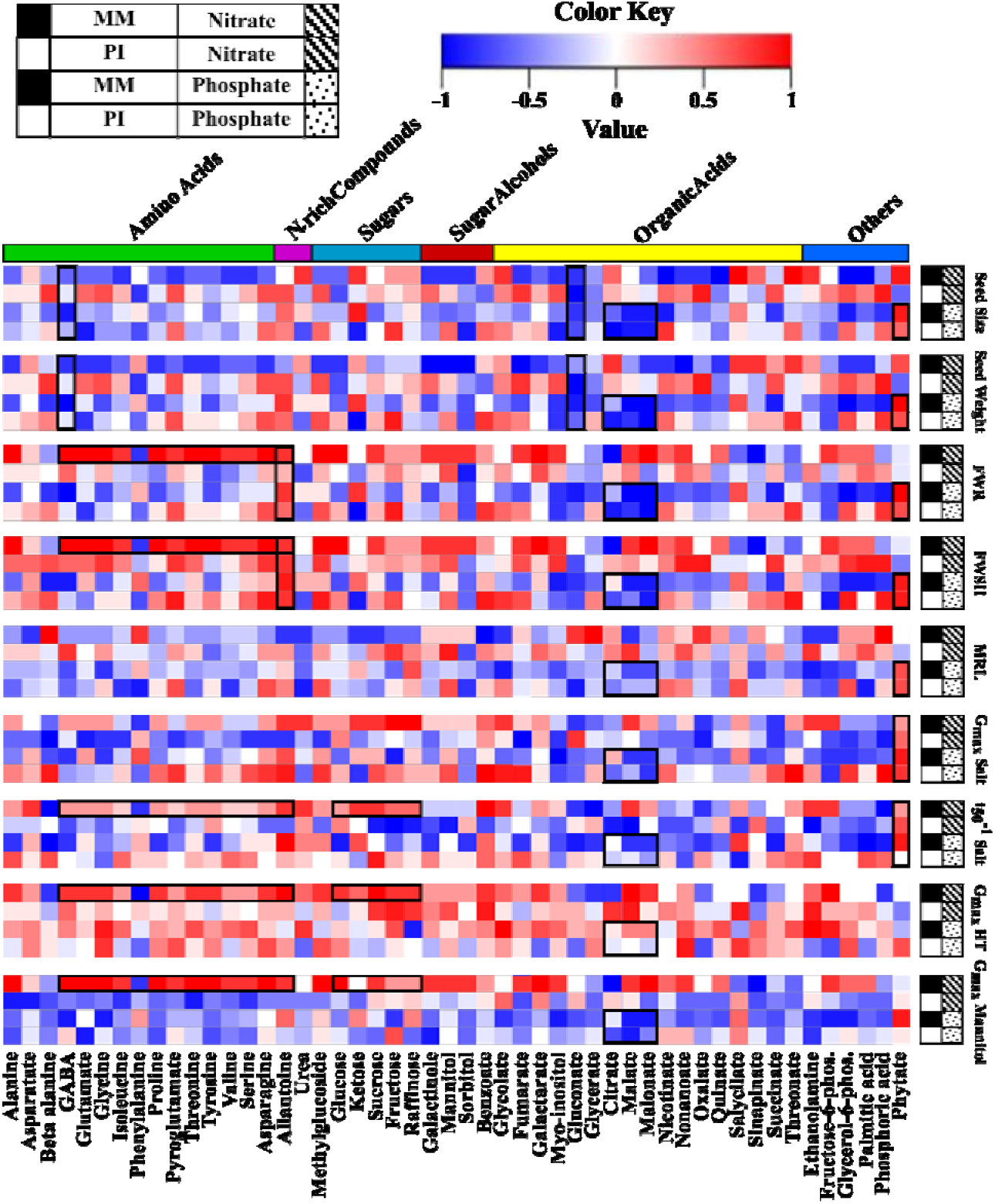
Correlation matrix of metabolites and seed and seedling quality traits. On right seed and seedling traits of two tomato genotypes: **MM** in black square and **PI** in white square in two different nutritional conditions: **Nitrate**, diagonal lines and **Phosphate**, dotted square are presented. At the bottom metabolites are presented in details and on top they are classified as groups of metabolites. Colour key table provides graphical representation of the correlation values of the traits and metabolites. The black rectangles indicate correlations mentioned in the result.

The correlation plots also indicated that some correlations were specific for either nitrate or phosphate environments. For seeds from plants grown in different phosphate environments, seed size and weight, FWR, FWSH and MRL of both genotypes displayed a negative correlation with some TCA cycle intermediates such as citrate, malate and malonate and positive correlation with phytate. Additionally, some correlations showed contrasting trends for the two environmental factors, such as the positive correlation of seed size and weight with citrate for seed batches originating from different maternal nitrate levels while they were negatively correlated in case of different phosphate levels (Fig. 7).

There are multiple correlations which were only present for a single condition. For instance in MM plants which were grown in different concentrations of nitrate, FWR and FWSH was positively correlated with a majority of the amino acids. In the same plants, seed germination quality traits under stressful conditions like salt, high temperature and osmotic stress were positively correlated with most amino acids and sugars (Fig. 7).

## Discussion

Although there are many reports addressing effects of the maternal environment on seed and seedling quality traits in several species, the effect of the maternal nutritional condition on seed performance has rarely been studied. In general, studying the effects of the maternal environment on seed performance may give insight into the processes that are involved in the adaptability of plants. The influence of the maternal environment on the next generation may be determined by several physiological traits such as germinability, size and weight of the seeds, as well as metabolic traits such as amino acid and sugar content of the seeds. Several studies have shown how the maternal environment affects seed and seedling quality in different crops. There are report on the effect of maternal photoperiod, temperature and nutrient conditions on seed performance (Demir *et al.*, 2004; Donohue, 2009; Munir *et al.*, 2001; Schmuths *et al.*, 2006). The influence of maternal nutrient conditions have been studied in different species such as tomato and Arabidopsis. It appeared that different dosages of maternal nutrients may influence seed characteristics such as seed size, seed weight and seed dormancy (Alboresi *et al.*, 2005; He *et al.*, 2014; Varis and George, 1985; Wulff, 1986). We here report the effect of a maternal environment with different concentrations of nitrate and phosphate on seed and seedling quality of two tomato genotypes. Additionally, we assessed primary metabolite profiles and analysed their correlation with different physiological traits such as seed germination and seedling development.

### Genotype by environment interactions (G×E)

We used two different genotypes to see how the nutritional maternal environment may influence seed and seedling quality traits and what is the effect of G×E interactions. We observed that the genotype is profoundly affecting seed and seedling characteristics and, thus, an obvious genotype specific effect was found for some phenotypic traits such as germination at high temperature (G_max_ HT) and metabolite content such as GABA (Table 2, Fig. 8). For the future, studying more *S. lycopersicum* genotypes may provide a more robust conclusion on the effect of the genotype. For phenotypic traits such as seed size and seed weight, there is a difference between the two genotypes, but there is no genotype specific effect since both genotypes are significantly influenced by nitrate and phosphate concentration (Fig. 8). MM and PI showed almost the same trend for traits such as seed size, seed weight, FWR, FWSH, DWR, DWSH and also MRL of the seedlings. However, MM plants produced generally bigger and heavier seeds and seedlings in all nutritional maternal environments (Fig. 2, Fig. 3). Furthermore, the highly significant influence of G×E interactions on several seed and seedling performance traits (Table 2) indicates that the phenotypic plasticity of the traits varied in relation to the different nutritional environments. In general, phosphate showed less effect than nitrate and among the nitrate levels, the traits were mostly influenced by 0N which could be an indication of the saturation of the nitrate response at the higher dosages in most of the traits (He *et al.*, 2014).

**Fig. 8.**
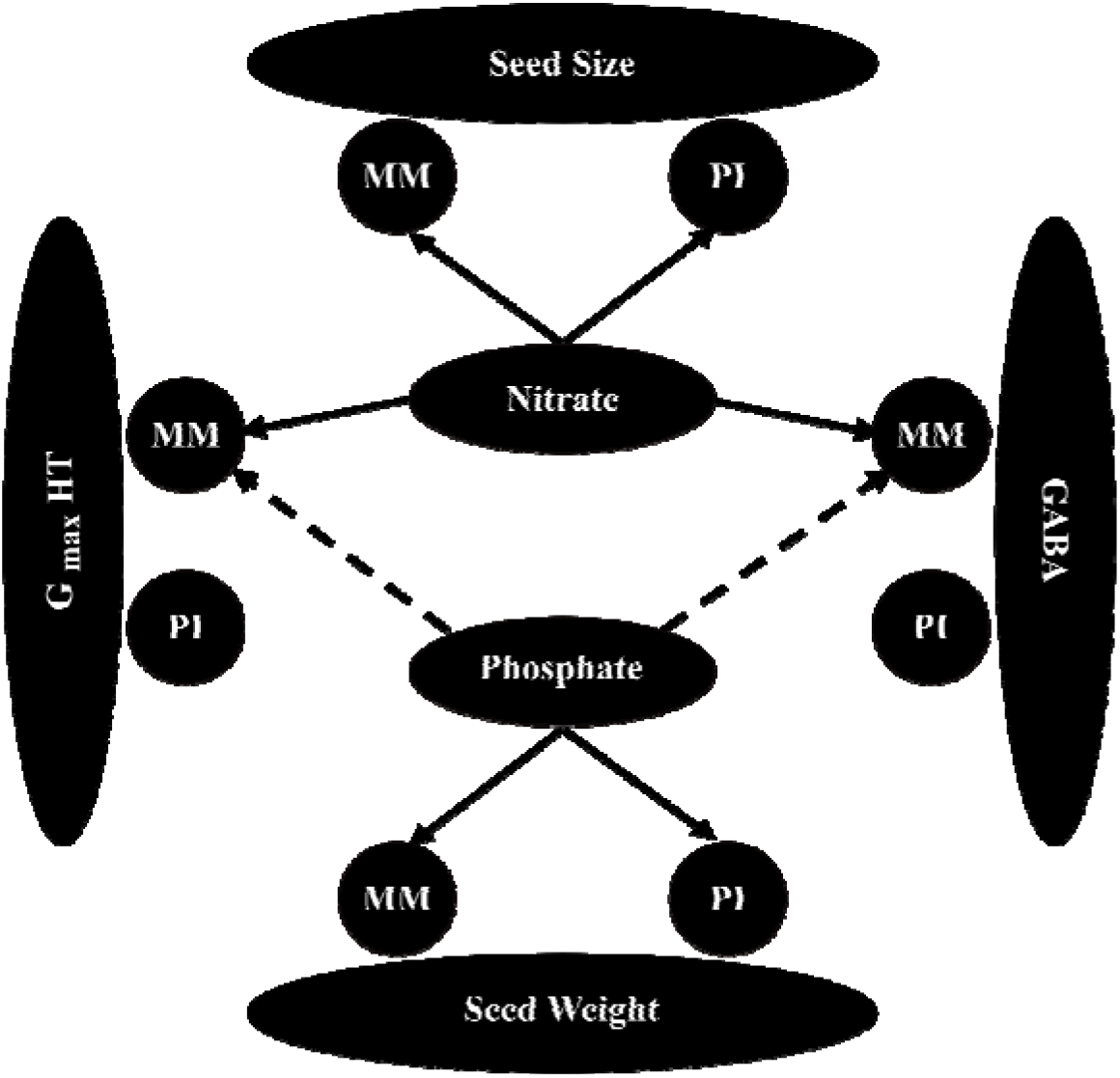
Summarizing the significant effects of nutritional maternal environments on seed size, seed weight, germination at high temperature (G_max_ HT) and the production of GABA in seeds. Nitrate positively regulated seed size for both genotypes while effect of nitrate on GABA accumulation and G_max_ HT was only observed when applied to MM seeds. Seed weight of both genotypes was positively regulated by phosphate content however, the (negative) effect of phosphate on GABA accumulation and G_max_ HT was only observed for MM seeds. Solid and dashed lines indicating the positive and negative effects, respectively.

### Relation between the nutritional environments of the mother plant and seed germination

There was also variability in the germination response of MM seeds from plants grown on different nitrate levels. Seeds that developed on higher levels of nitrate germinated better under stressful conditions, such as osmotic, salt and high temperature. These seeds also contained higher contents of amino acids. Several studies have implied that in response to stressful conditions, amino acids can be fed into the TCA cycle and serve as the main substrate for energy generation. This might explain higher seed germination percentages under stress conditions (Galili, 2011). Although different concentrations of nitrate and phosphate altered seed germination percentages under stressful conditions in MM, there was no significant change of seed germination in PI since PI showed almost 100% germination under optimal and the tested stressful germination conditions. PI is a wild tomato species and is often more tolerant to various biotic and abiotic stresses (Kazmi *et al.*, 2012; Kumar, 2006; Rao *et al.*, 2013; Rodríguez-López *et al.*, 2011). Loss of abiotic stress tolerance in tomato cultivars is thought to be the result of genetic bottlenecks during domestication (Bai and Lindhout, 2007; Doebley *et al.*, 2006).

### Seed size and seedling growth are strongly influenced by the maternal nutritional environment

As described above, it appears that for both genotypes increasing the nitrate level leads to higher amounts of amino acids in the seeds. Furthermore, proteins are one of the principal storage compounds of seeds that are subsequently used as nutrients and energy source to assert seed germination and seedling establishment (Bewley *et al.*, 2012; Galili *et al.*, 2015). Thus, the higher dosage of nitrate may result in the synthesis of more amino acids during development and this might increase protein content which subsequently might result in bigger and heavier seed and seedling production and eventually successful establishment of seedlings (Castro *et al.*, 2006; Ellis, 1992). Seedling vigour and establishment are two essential parameters that may influence final crop yield and are therefore necessary for profitable crop production. Successful seedling establishment can be considered as the most critical stage of crop development. Such an important stage can be influenced by parameters such as the maternal environment in which the seeds mature and several seed characteristics such as seed size, seed weight and stored organic and mineral nutrients in the produced seeds (Lamont and Groom, 2013; Stevens *et al.*, 2014). Confirming a study by Khan *et al.* (2012), we show that seedling size in tomato is positively correlated with seed size and weight in both genotypes. The positive correlation that we found between seed and seedling size is also in agreement with several other studies (Cornelissen, 1999; Greene and Johnson, 1998; Khan *et al.*, 2012). Additionally, we have shown that increasing the maternal phosphate level enhanced seed size and seed weight which again resulted in increased seedling size. We observed that higher amounts of phosphate decreased the amount of F6P in seeds. Moreover, the level of citrate and malate in the seeds decreased with increasing maternal phosphate levels. Since glycolysis and the TCA cycle are key metabolic pathways by which organisms generate energy, decrease in the level of intermediates of these pathways like F6P, citrate and malate possibly indicates their consumption for energy production. It might suggest that higher utilization of glycolytic and TCA intermediates in seeds of higher maternal phosphate concentrations, results in more production of ATP and consequently more growth of the seedlings (Fig. 3, Fig. 6). In addition, production of bigger and heavier seedling for seeds developed under higher dosage of phosphate may be related to the higher amount of reserves which could be stored in bigger tomato seeds produced under the same condition.

### Role of GABA in plant adaptation

Carbon (C) and nitrogen (N) are two vital factors that help plants to execute essential cellular activities. C and N metabolic pathways are strongly coordinated to ensure optimal growth and development in plants (Zheng, 2009). Several studies have reported that when plants are facing N deficiency, photosynthetic output and, consequently, plant growth is negatively influenced (Coruzzi and Bush, 2001; Coruzzi and Zhou, 2001). Several studies have implicated a primary role of the GABA shunt in the central C/N metabolism (Fait *et al.*, 2011). In this study we found that the application of lower amounts of nitrate to mother plants resulted in lower production of GABA in the seeds of the progeny. Therefore, the decrease in GABA content in dry seeds as a result of maternal N deficiency could be an indication of GABA usage to alleviate N shortage and, subsequently, to recover the C/N balance (He *et al.*, 2016).

In this study, we observed the highest percentage of seed germination under high temperature conditions in seeds that had developed in high levels of nitrate and/or low levels of phosphate (Fig. 1). On the other hand, although enhancing the maternal nitrate level results in an increase in the GABA content of the seeds of MM, enhancing the phosphate levels conversely decreased it (Fig. 6). Thus, there is a good correlation for MM seeds between the different GABA levels in the seeds as a result of the maternal environment and the ability to germinate at high temperatures. This is in agreement with many studies in which GABA has been shown to act as an abiotic stress mitigating component in plants (Bouche and Fromm, 2004; Kinnersley and Turano, 2000).

In this study we observed that different dosages of nitrate and phosphate during seed development and maturation may influence the seed and seedling characteristics. We have shown that in tomato, nitrate has a greater effect on seed and seedling performance as compared with phosphate. However, two different tomato genotypes showed different responses to the maternal environment and sometimes genetic specific responses were observed for some traits. Such differential responses may indicate the contribution of different genetic and molecular pathways to the phenotypic adaptation. Further investigating such observations as well as the effect of G×E interaction on the performance of the tomato seed and seedling may ultimately help in predicting and improving seed and seedling quality by controlling production environments and breeding programs.

## Supplementary data

**Table S1.** List of the nutrient solutions with their concentrations used for different growing environments of tomato plants.

**Supplementary Excel File S1.** Detected metabolites in the seeds of two tomato genotypes (**MM** and **PI**) grown in different concentration of nitrate and phosphate.

**Figure S1.** Effects of maternal nutritional environments on seed germination traits of both **MM** and **PI**.

**Figure S2.** Effects of maternal nutritional environments on ABA **(A)** and Phytate **(B)** content of both **MM** and **PI** seeds developed in different concentrations of nitrate and phosphate.

**Figure S3.** Effects of maternal nutritional environments on seedling quality traits of both **MM** and **PI**.

**Figure S4.** Cluster analysis of known primary metabolites in **MM** and **PI** seeds in response to different concentration of nitrate and phosphate during maternal growth.

## Acknowledgments

This work was supported by Technology Foundation (STW), which is part of the Netherlands Organization for Scientific Research (NWO) (NG, LW, WL) and by the Iranian Ministry of Science, Research and Technology (SS).

## Abbreviations

AUC: area under the germination curve
DWR: dry weight of root
DWSH: dry weight of shoot
FWR: fresh weight of root
FWSH: fresh weight of shoot
G_max_: maximum germination
G×E: genotype by environment interactions
HCL: hydrochloric acid
HT: high temperature
MM: *Solanum lycopersicum* cv. Money maker
MRL: main root length
NLR: number of lateral root
PCA: principle component analysis
PI: *Solanum pimpinellifolium*
T_50_^-1^: reciprocal of time to reach 50% of germination
TCA: tricarboxylic acid
U_8416_: uniformity of germination or time from 16% till 84% germination

